# Widespread Neuroanatomical Integration and Distinct Electrophysiological Properties of Glioma-Innervating Neurons

**DOI:** 10.1101/2024.08.25.609573

**Authors:** Annie L. Hsieh, Sanika Ganesh, Tomasz Kula, Madiha Irshad, Emily Anne Ferenczi, Wengang Wang, Yi-Ching Chen, Song-Hua Hu, Zongyu Li, Shakchhi Joshi, Marcia C. Haigis, Bernardo L. Sabatini

## Abstract

Gliomas are the most common malignant primary brain tumors and are often associated with severe neurological deficits and mortality. Unlike many cancers, gliomas rarely metastasize outside the brain, indicating a possible dependency on unique features of brain microenvironment. Synapses between neurons and glioma cells exist, suggesting that glioma cells rely on neuronal inputs and synaptic signaling for proliferation. Yet, the locations and properties of neurons that innervate gliomas have remained elusive. In this study, we utilized transsynaptic tracing with a pseudotyped, glycoprotein-deleted rabies virus to specifically infect TVA and glycoprotein-expressing human glioblastoma cells in an orthotopic xenograft mouse model, allowing us to identify the neurons that form synapses onto the gliomas. Comprehensive whole-brain mapping revealed that these glioma-innervating neurons (GINs) consistently arise at brain regions, including diverse neuromodulatory centers and specific cortical layers, known to project to the glioma locations. Molecular profiling revealed that these long-range cortical GINs are predominantly glutamatergic, and subsets express both glutamatergic and GABAergic markers, whereas local striatal GINs are largely GABAergic. Electrophysiological studies demonstrated that while GINs share passive intrinsic properties with cortex-innervating neurons, their action potential waveforms are altered. Our study introduces a novel method for identifying and mapping GINs and reveals their consistent integration into existing location-dependent neuronal network involving diverse neurotransmitters and neuromodulators. The observed intrinsic electrophysiological differences in GINs lay the groundwork for future investigations into how these alterations may correspond with the postsynaptic characteristics of glioma cells.

**Significance:** We have developed a novel system utilizing rabies virus-based monosynaptic tracing to directly visualize neurons that synapse onto human glioma cells implanted in mouse brain. This approach enables the mapping and quantitative analysis of these glioma-innervating neurons (GINs) in the entire mouse brain and overcomes previous barriers of molecular and electrophysiological analysis of these neurons due to the inability to identify them. Our findings indicate that GINs integrate into existing neural networks in a location-specific manner. Long-range GINs are mostly glutamatergic, with a subset expressing both glutamatergic and GABAergic markers and local striatal GINs are GABAergic, highlighting a complex neuromodulatory profile. Additionally, GINs exhibit unique action potential characteristics, distinct from similarly selected neurons in non-tumor-bearing brains. This study provides new insights into neuronal adaptations in response to forming putative synapses onto glioma, elucidating the intricate synaptic relationship between GINs and gliomas.

## Introduction

Neuronal interactions with glioma cells are critical for glioma growth (1, 2). These interactions are characterized by the formation of synapse-like structures between neurons and glioma cells, whose electrical stimulation elicits calcium influx and post-synaptic currents in glioma cells (3–5). Neuronal activity promotes glioma tumorigenesis by secretion of synaptic protein neuroligin 3 and brain-derived neurotrophic factor (BDNF) and enhances invasive behavior via axon guidance protein SEMA4F (6–9). Furthermore, gliomas can remodel human neural circuits, leading to neuronal hyperexcitability and altered functional connectivity, thereby adversely impacting patient survival (10). Previous studies have implicated glutamatergic signaling, mediated through AMPA-type glutamate receptors (AMPARs), as the primary mechanism of neuron-glioma interactions (3, 4).

Despite these advances, the specific neuronal populations that innervate gliomas remain poorly defined. Transsynaptic tracing using rabies virus has significantly enhanced our understanding of neuronal circuits by enabling detailed mapping of neurons that are putatively presynaptic to specific brain regions or genetically-defined cells (11–13). Similar tracing techniques have been used to study interactions between neurons and oligodendrocyte precursor cells (14). The introduction of the N2c rabies virus strain has further improved the efficiency of transsynaptic mapping and prolonged the viability of rabies virus-infected neurons (15).

In this study, we map and profile glioma-innervating neurons (GINs) using rabies virus-based transsynaptic tracing in human glioma orthotopic xenograft mouse models. This approach enables brain-wide mapping of neuronal networks interacting with glioma cells and reveals a consistent pattern that is dependent on glioma location. The anatomical distribution of GINs closely mirrors that of neurons that typically innervate the affected regions, indicating that gliomas synaptically integrate into pre-existing neural circuits. Fluorescence in situ hybridization (FISH) revealed that most long-range GINs are glutamatergic, with a subset also displaying both glutamatergic and GABAergic markers, whereas local inputs to striatal glioma are GABAergic. Electrophysiology revealed that GINs exhibit distinct action potential waveforms compared to those of neurons that normally innervated the tumor location. In summary, we demonstrate that RV-mediated trans-synaptic tracing can be used to reveal the network that is presynaptic to gliomas, permitting future analysis of the molecular and physiological properties of such networks.

## Results

### Brain-wide distribution of glioma-innervating neurons revealed by rabies virus-based transsynaptic tracing

To determine the anatomical locations of putative glioma-innervating neurons (GINs), we adapted a rabies virus (RV)-based transsynaptic tracing approach (16). We engineered the human glioblastoma cell line LN229 to stably express the avian sarcoma leucosis virus subgroup A receptor (TVA) and green fluorescent proteins (eGFP), with or without the rabies glycoprotein (N2cG) (Fig. 1*A*, Fig. S1*A*). We sorted eGFP-positive cells to obtain pure populations of TVA-eGFP (Ctrl) and TVA-N2cG-eGFP (G-complement) cells (Fig. 1*A*, Fig S1*B*). Expression of TVA, an avian receptor protein that is absent in mammalian cells, in the Ctrl or G-complemented LN229 cells allows the entry of RVs pseudotyped with the avian sarcoma leucosis virus glycoprotein type A (EnvA) (16). Transsynaptic spread to presynaptic partners of the G-complemented LN229 cells is enabled by the complementation of the glycoprotein gene-deleted RVs by N2cG. Lack of TVA in neighboring neurons and astrocytes prevents direct RV infection of these cells whereas lack of G expression in any neurons that RV enters by trans-synaptic spread prevents further movement beyond these neurons that directly contact glioma cells. A reporter gene (red-fluorescent protein, mCherry) encoded in the RV genome labels the synaptically connected first-degree neurons and the primarily infected LN229 cells, which also express GFP.

**Figure. 1.**
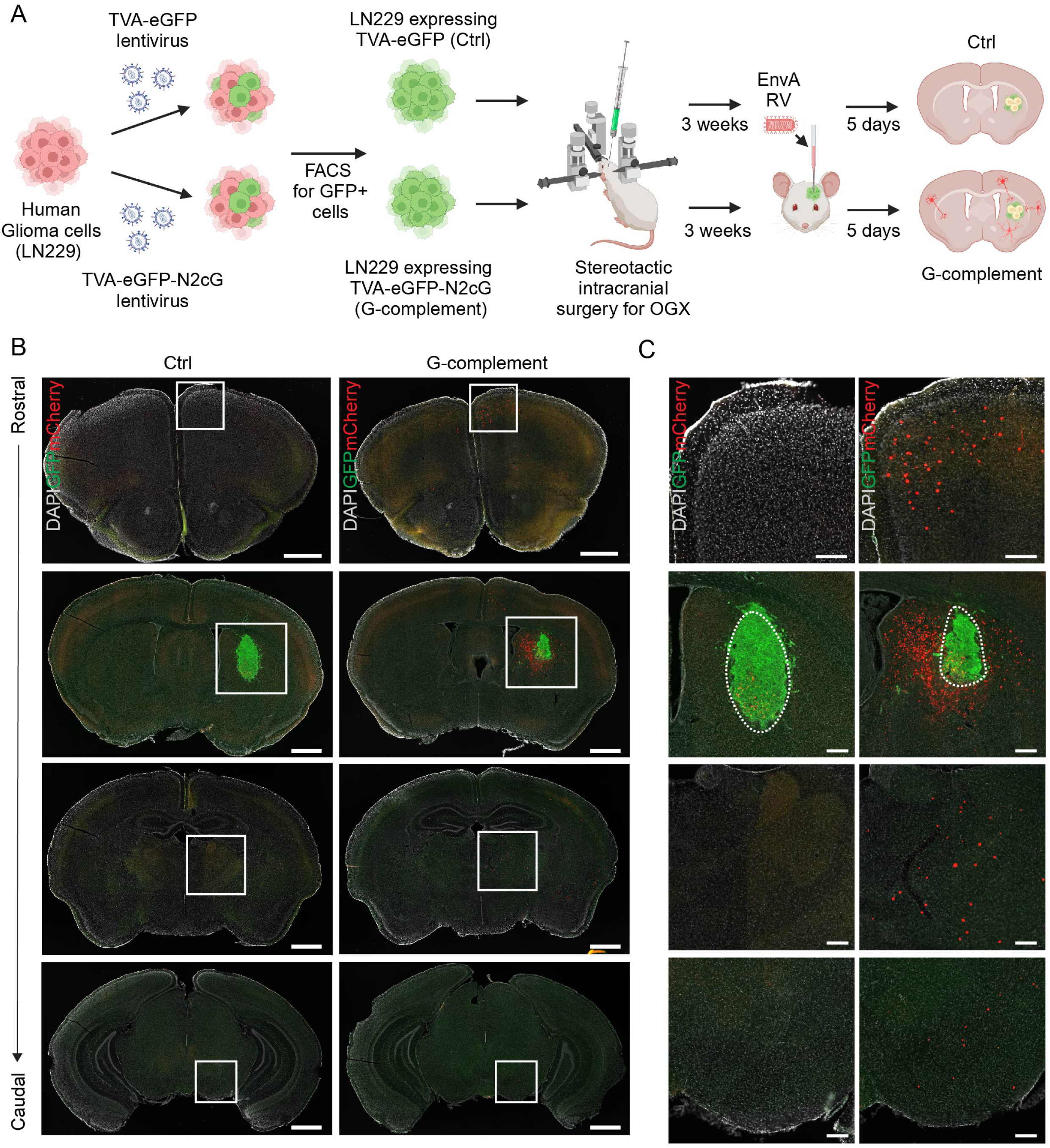
Transsynaptic tracing of glioma-innervating neurons using rabies virus. (A) Schematic of the experimental workflow used to establish stable human glioma Ctrl (TVA-eGFP) and G-complement (TVA-eGFP-N2cG) LN229 orthotopic glioma xenograft (OGX). This is followed by the injection of pseudotyped N2cG-H2b-mCherry rabies virus (EnvA RV) for monosynaptic retrograde transsynaptic tracing of glioma-innervating neurons. FACS: Fluorescence-activated cell sorting. Schema was created with BioRender.com. (B) Representative coronal brain sections from Ctrl and G-complement striatal OGX following RV-based tracing. Nuclear-localized RV H2b-mCherry (red) specifically labels neurons in G-complement OGX implanted brains but not Ctrl OGX implanted brains. Scale bar: 1mm. (C) Inset from panel (B) showing detailed localization of labeled neurons. Scale bar: 100µm.

Ctrl and G-complement LN229 cells were orthotopically injected into the striatum of immunocompromised NCG mice to establish orthotopic glioma xenografts (OGX). After three weeks to allow for the glioma cells to expand and potentially integrate with host circuits, we injected pseudotyped (EnvA), N2cG-deleted, mCherry-expressing RV into the Ctrl- and G-complement glioma xenograft at identical stereotaxic coordinates. The RV specifically targeted TVA-expressing LN229 cells via EnvA-TVA interaction and, when complemented by glioma-expressed G-protein, traverse trans-synaptically to presynaptic neurons connected to the infected glioma cells. We visualized RV-N2cG-mCherry-labeled cells five days post-injection using fluorescence microscopy (Fig. 1*B* and *C*). Immunohistochemical analysis confirmed that all mCherry-positive cells were NeuN positive and GFAP negative, confirming their neuronal identity (Fig. S1*D*). These neurons were found both locally in the striatum surrounding the tumor as well as distant from the tumor injection site, demonstrating the extensive transsynaptic spread of the virus (Fig. 1*B* and *C*). mCherry-positive neurons were observed exclusively in G-complement OGX-harboring mice. In contrast, RV injection into Ctrl OGX-harboring mice resulted in robust starter LN229 cell labeling but no trans-synaptic labeling to neurons, underscoring the specificity of this approach in identifying GINs.

### Glioma integrates into existing neuronal networks in a location-dependent manner

Human gliomas arise in various brain regions, necessitating modeling that reflects their diverse microenvironment (17). We selected specific brain regions for glioma cell injections based on their typical occurrence sites and distinct neuronal connectivity. To determine how glioma locations influence neuronal projections and assess if GINs maintain consistent features across different brain areas, we compared GIN distributions for OGX tumors implanted in the striatum, thalamus and motor cortex, each of which possesses distinct circuitry and neuromodulatory pathways (Fig. 2*A*).

**Figure. 2.**
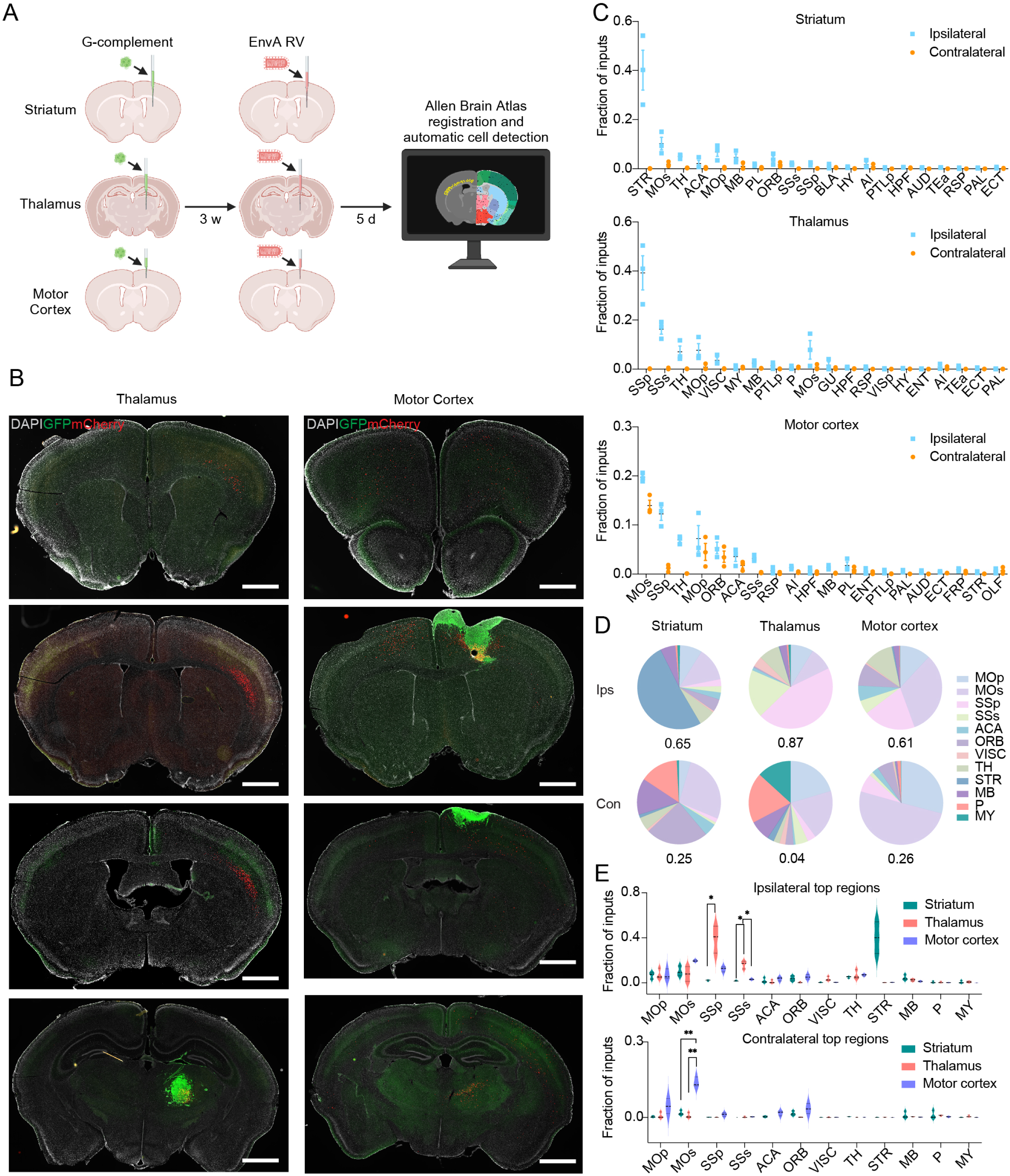
Comprehensive brain-wide mapping of glioma-innervating neurons (GINs). (A) Schematic diagram illustrating the experimental workflow and analysis pipeline for assessing the location-dependent network of GINs. G-complement: TVA-eGFP-N2cG LN229 human glioma cells; EnvA RV: pseudotyped N2cG-H2b-mCherry rabies virus; 3 w: 3 weeks; 5 d: 5 days. (B) Representative coronal brain sections from LN229 G-complement orthotopic glioma xenograft implanted in the thalamus and motor cortex, followed by pseudotyped N2cG-H2b-mCherry rabies virus for monosynaptic retrograde transsynaptic tracing of glioma-innervating neurons. Scale bar: 1mm. (C) Top 20 input regions to striatal, thalamic and motor cortex LN229 G-complement OGX, presenting the distribution of labeled neurons ipsilateral and contralateral to the glioma. Data collected from three mice per glioma location (n=3). Brain regions are abbreviated as follows: STR, striatum; MOs, secondary motor area; TH, thalamus; ACA, anterior cingulate area; MOp, primary motor area; MB, midbrain; PL, prelimbic area; ORB, orbital area; SSs, supplemental somatosensory area; SSp, primary somatosensory area; BLA, basolateral amygdalar nucleus; HY, hypothalamus; AI, agranular insular area; PTLp, posterior parietal association areas; HPF, hippocampal formation; AUD, auditory areas; TEa, temporal association areas; RSP, retrosplenial area; PAL, pallidum; ECT, ectorhinal area; VISC, visceral area; MY, medulla; P, pons; GU, gustatory areas; VISp, primary visual area; ENT, entorhinal area; FRP, frontal pole/ cerebral cortex; OLF, olfactory areas. (D, E) Distribution of GINs across the top 12 regions among all three OGX sites, illustrated in pie charts (D) and violin plots (E) to depict the extent of neuronal involvement ipsilateral (Ips) and contralateral (Con) to the glioma. Number below the pie chart represents the total fraction. Statistically significance denoted by asterisks. \**P* < 0.05; \*\**P* < 0.01; multiple unpaired t-tests, adjusted for multiple comparisons using the Holm-Šídák method.

Following RV injection into G-complement OGX tumors in the thalamus or motor cortex, mCherry-positive neurons indicative of GIN connections were detected across multiple brain areas. In contrast, no mCherry-positive cells were seen in Ctrl OGX animals, except within the starter cell population (Fig. 2*B* and Fig. S1*C*). GIN distribution varied significantly depending on the tumor’s location: striatal gliomas primarily connected with local and ipsilateral motor cortical neurons; thalamic gliomas received projections from deep cortical layers of the ipsilateral somatosensory cortex; motor cortex gliomas had inputs from bilateral prefrontal, motor and somatosensory cortices, as well as the ipsilateral thalamus (Fig. 1*B* and *C*, Fig. 2*B*).

To quantitatively compare GINs across different glioma locations, we utilized whole-brain imaging and automated analysis. We employed an RV expressing nuclear-localized mCherry (CVS-N2c-dG-H2b-mCherry) to facilitate precise counting of rabies-infected neurons and measurement of the distribution of putative neuronal inputs to gliomas. Whole brain fluorescence image sections were analyzed by NeuroInfo software (MBF Bioscience) for automatic cell detection and alignment to the Allen Brain Atlas for comparison across different animals (Fig. 2*A*). The fraction of H2b-mCherry-positive neurons in each brain region defined by Allen Brain Atlas was calculated to quantify GIN distribution (Fig. 2*C* and *D*). In the striatal glioma group (n=3), the majority of such cells were found in the ipsilateral hemisphere including ∼40% of all labeled cells in striatum, ∼10% in secondary motor area, 6.8% in primary motor cortex (Fig. 2*C* and *D*; STR 40.2 ± 8%, MOs 10.2 ± 2.6%, MOp 6.8 ± 1.8% of total inputs). Most of the contralateral GINs were located in the secondary motor area (1.8% of all cells) and orbital area (1.6%) (Fig. 2*C* and *D*; MOs 1.8 ± 0.56%, ORB 1.6 ± 0.5% of total inputs). The thalamic glioma group (n=3) showed 39% of GINs in the ipsilateral primary somatosensory area, 16% in the supplemental somatosensory area, and 8% in the secondary motor area (Fig. 2*C* and *D*; SSp 39.3 ± 7%, SSs 16.3 ± 2%, MOs 7.9 ± 3.8% of the total inputs), with the largest fraction of contralateral inputs from the primary motor area (0.8%), secondary motor area (0.77%) and pons (0.77%) (Fig. 2*C* and *D*; MOp 0.8 ± 0.7%, MOs 0.77 ± 0.6%, P 0.77 ± 0.1% of the total inputs). In the motor cortex glioma group (n=3), 20% of ipsilateral GINs and 14% of contralateral GINs project from the secondary motor area, 12% from the ipsilateral primary somatosensory area, and 7% from the ipsilateral primary motor area and thalamus (Fig. 2*C* and *D*; MOs (ipsilateral) 19.7 ± 0.5%, MOs (contralateral) 13.9 ± 1%, MOp 7.2 ± 2.6%, TH 7 ± 0.5% of the total inputs). Comparing GIN distribution between striatal, thalamic and motor cortex glioma groups, GIN fraction is significantly higher in ipsilateral primary and supplementary somatosensory area in thalamic glioma group than striatal glioma group (Fig. 2E). Contralaterally, GIN fraction is significantly higher in secondary motor cortex in motor cortex glioma groups than both striatal and thalamic glioma groups (Fig. 2E).

### Cortical layer specificity and neuromodulatory pathways differ across glioma locations

We observed variability in the cortical layers of RV-labeled putative GINs depending on tumor locations: cortical neurons projecting to striatal gliomas predominantly originated from layer 5 bilaterally (66.1% ± 2.5% of total isocortex inputs ipsilaterally, 83.5% ± 3.3% contralaterally), whereas those projecting to thalamic gliomas mainly came from layer 6 bilaterally (92% ± 2.6% ipsilaterally, 73.8% ± 11% contralaterally) (Fig. 3*A* and *B*). Interestingly, cortical neurons projecting to motor cortex gliomas are located in both layers 5 (37.2% ± 4.7%) and 6 (47.6% ± 5.6%) ipsilaterally (Fig. 3A and B), suggesting that layer 6 corticocortical neurons, which typically extend long horizontal axons across cortical areas, form synapses with gliomas (18). To investigate the potential type of neurotransmitters or neuromodulators used by GINs, we analyzed the whole-brain rabies mapping dataset, mapping GINs to brain regions identified in previous literature as being involved in location-specific neuromodulatory centers (19–21). This analysis revealed that putative GINs are found in centers that release a diverse array of neurotransmitters and neuromodulators. For instance, striatal gliomas receive inputs from glutamatergic centers (cortex, thalamus (TH), anterior cingulate area (ACA), basolateral amygdalar nucleus (BLA) and hippocampal formation (HPF)), dopaminergic centers (ipsilateral substantia nigra/compact part (SNc) and ventral tegmental area (VTA)) and cholinergic centers (bilateral pedunculopontine nucleus (PPN)). Thalamic gliomas receive inputs from glutamatergic centers in the cortex, histaminergic centers in the hypothalamus (HY), cholinergic centers in the basal forebrain (BF) and pedunculopontine nucleus (PPN), dopaminergic centers from ipsilateral zona incerta (ZI) (22), bilateral periaqueductal gray (PAG) (23) and bilateral parabrachial nucleus (PB) (24), as well as GABAergic centers including ipsilateral ZI, substantia nigra/ reticular part (SNr) and globus pallidus externa (GPe). Motor cortex gliomas receive inputs from glutamatergic centers (cortex and TH), cholinergic centers (BF), GABAergic center (ipsilateral GPe/GPi), dopaminergic centers (ipsilateral VTA) and serotonergic centers (dorsal raphe nucleus (DR)) (Fig. 3*C*). These findings suggest that gliomas intergrade into existing neuronal networks in a location-dependent manner, engaging a variety of neuromodulatory inputs.

**Figure. 3.**
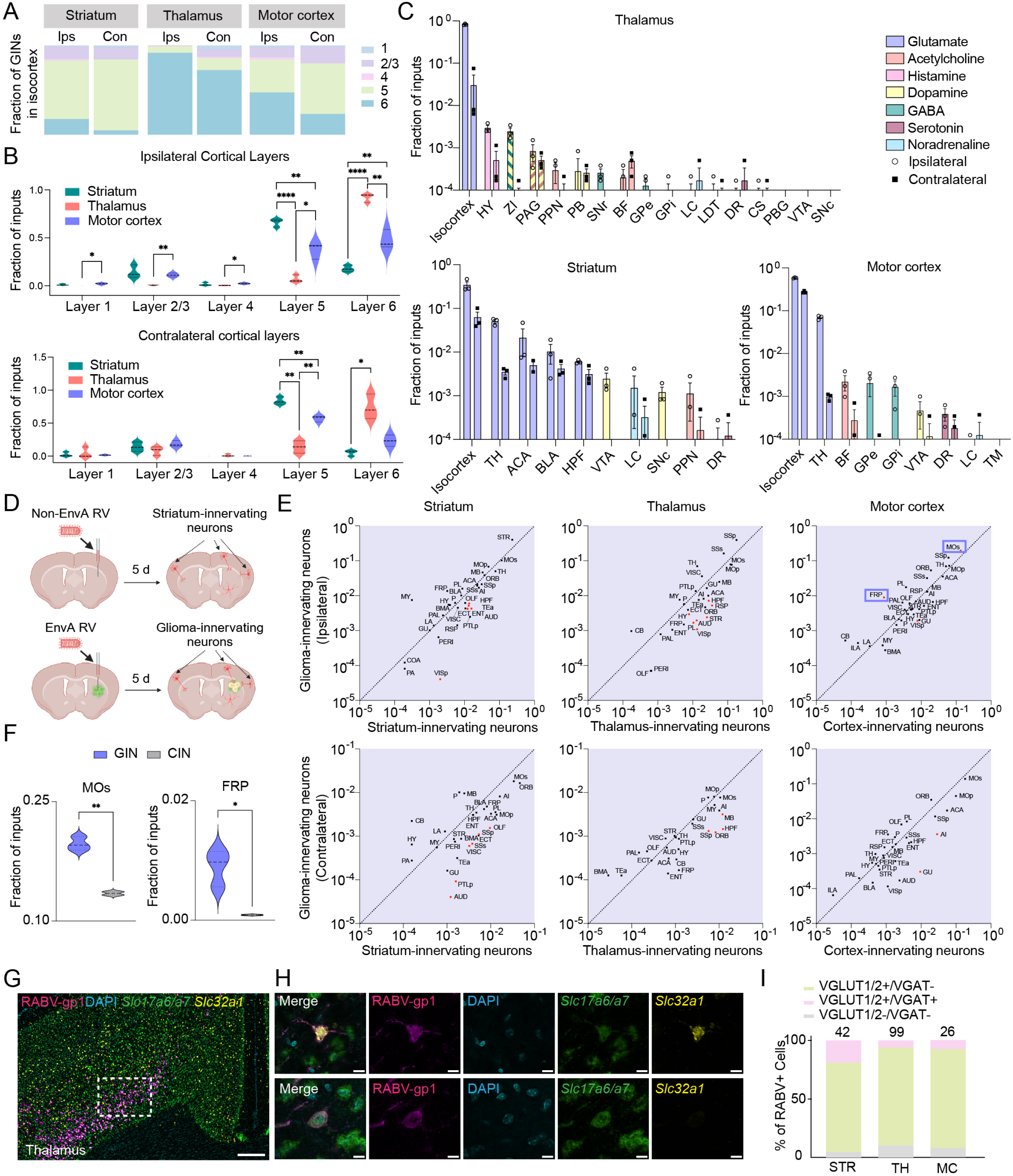
Glioma-innervating neurons (GINs) exhibit diverse neuromodulatory pathways. (A, B) Distribution of GINs across cortical layers, illustrated with pie charts (A) and violin plots (B) to demonstrate cortical layer specificity in a glioma location-dependent manner. (Ips, ipsilateral to glioma; Con, contralateral to glioma; 1, layer 1; 2/3, layer 2/3; 4, layer 4; 5, layer 5; 6, layer 6). Statistically significance denoted by \**P* < 0.05; \*\**P* < 0.01, \*\*\*\**P* < 0.0001; multiple unpaired t-tests, adjusted for multiple comparisons using the Holm-Šídák method. (C) Mapping of GINs in key neuromodulatory brain regions related to striatal, thalamic and motor cortex LN229 G-complement OGX, showing labeled neurons ipsilateral and contralateral to the glioma (n=3 mice for striatal, thalamus and motor cortex OGX). HY, hypothalamus; ZI, zona incerta; PAG, Periaqueductal gray; PPN, Pedunculopontine nucleus; PB, Parabrachial nucleus; SNr, substantia nigra/ reticular part; BF, basal forebrain (including striatum ventral region, pallidum/ ventral region, pallidum/ medial region,pallidum/ caudal region); GPe, globus pallidus/ external segment; GPi, globus pallidus/ internal segment; LC, locus ceruleus; LDT, laterodorsal tegmental nucleus; DR, dorsal nucleus raphe; CS, superior central nucleus raphe; PBG, parabigeminal nucleus; VTA, ventral tegmental area; SNc, substantia nigra/ compact part; TM, Tuberomammillary nucleus. (D) Schematic illustrating the experimental design for assessing the distribution of GINs relative to physiological inputs in the striatum. Non-EnvA RV: nonpseudotyped CVS-N2c-dG-TdTomato rabies virus; EnvA RV: pseudotyped N2cG-H2b-mCherry rabies virus; 5 d: 5 days. (E) Scatter plots illustrating the distribution of GINs from striatal, thalamic and motor cortex OGX to their respective region-innervating neurons. Each dot represents the average fraction of inputs at each brain region with red dots indicating statistically significant differences in the fraction of GINs versus physiological inputs. Analysis was performed using multiple unpaired t-tests, not adjusted for multiple comparisons. (F) Violin plots showing the fraction of GINs in the ipsilateral secondary motor area (MOs) and frontal pole/cerebral cortex (FRP) compared to that of cortex-innervating neurons (CINs) in the same area. \**P* < 0.05; \*\**P* < 0.01, unpaired t-tests. (G) Representative in situ hybridization confocal image from the ipsilateral motor cortex of thalamic OGX. RABV-gp1(rabies virus complete genome)-positive cells were classified as VGLUT1/2+ (*Slc17a6* or *Slc17a7 positive*)/VGAT+(*Slc32a1 positive*), VGLUT1/2+/VGAT- or VGLUT1/2-/VGAT-(no VGLUT1/2-/VGAT+ cells observed). Scale bar: 250μm. (H) Representative in situ hybridization confocal image showing a VGLUT1/2+/VGAT+ cell (top) and VGLUT1/2+/VGAT-cell (bottom). Scale bar: 10 μm. (I) Bar graph quantifying the percentages of RABV+ cells that are VGLUT1/2+/VGAT+, VGLUT1/2+/VGAT- or VGLUT1/2-/VGAT-in striatal (STR), thalamic (TH) and motor cortex (MC) OGX animals (n=1 mouse per glioma location).

### Upstream neuronal networks projecting into gliomas reflect physiological networks

Although the distribution of GINs largely aligns with the expected neuronal inputs for each glioma location, it remains uncertain whether gliomas exhibit a preference for innervation by specific neuronal pathways. To assess potential differences between the distribution of GINs and of the input neurons that normally innervate each brain area in the absence of a tumor, we injected a non-pseudotyped, glycoprotein-deleted rabies virus (CVS-N2c-dG-TdTomato) at the coordinates corresponding to the glioma sites (Fig. 3*D* and S2*A*). This virus will directly infect synaptic terminals but not move trans-synaptically. Whole-brain sections were analyzed using the same Neuroinfo pipeline to quantitatively compare regions containing GINs and the native input neurons with different glioma locations (Fig. 3*D* and S2*B*). Across all 39 brain regions examined, no statistically significant differences (multiple unpaired t-tests, adjusted for multiple comparisons using the Holm-Šídák method) in the fraction of total inputs between GINs and the native input neurons were observed, indicating that the distribution of GINs mirrors that of neurons physiologically projecting to the glioma sites (Fig. 3*D*).

Further analysis of the number of labeled neurons in individual brain regions revealed greater labeling by direct terminal infecting RV than by the trans-synaptic labeling from GINs, likely a result of more efficient retrograde tracing by the nonpseudotyped, glycoprotein-deleted rabies virus (Fig. 3*E*, red dots). For instance, compared to striatum-innervating neurons, fewer GINs were observed in regions typically interacting with the striatum physiologically, such as ipsilateral primary visual area (VISp), hippocampal formation (HPF), temporal association areas (TEa), auditory areas (AUD), entorhinal area (ENT), and contralateral auditory areas (AUD), supplemental somatosensory area (SSs), olfactory areas (OLF), posterior parietal association areas (PTLp), primary somatosensory area (SSp), and visceral area (VISC) (Fig. 3*E*, red dots). Similarly, compared with thalamic-innervating neurons, fewer GINs were observed in ipsilateral striatum (STR), auditory areas (AUD), primary visual area (VISp), prelimbic area (PL), hypothalamus (HY), retrosplenial area (RSP), hippocampal formation (HPF), and contralateral midbrain (MB), hippocampal formation (HPF), orbital area (ORB), primary somatosensory area (SSp) (Fig. 3*E*, red dots). In contrast, GINs in motor cortex gliomas closely aligned with the distribution of cortex-innervating neurons, with notable exceptions including more GINs in ipsilateral secondary motor area (MOs) and frontal pole/cerebral cortex (FRP), and fewer in ipsilateral primary visual area (VISp), contralateral agranular insular area (AI) and gustatory areas (GU) (Fig. 3*E*). The significantly increased fraction of GINs in ipsilateral MOs and FRP, both largely glutamatergic areas, suggests that glioma may preferentially connect with the synaptic terminals of neurons from these regions (Fig. 3*F*).

### Molecular characterization reveals the diverse neurotransmitter signatures of glioma-innervating neurons

Single-cell sequencing of human glioma tissues uncovered an enrichment of AMPA receptors in gliomas compared to normal neighboring cells, supporting the hypothesis that GINs are glutamatergic (3, 4). Our analysis of cortical layers and neuromodulatory pathways also demonstrated that GINs consistently cluster in areas rich in glutamatergic neurons, irrespective of glioma location (Fig. 3*A, B* and *C*). To further characterize the molecular properties of GINs, we conducted mRNA-fluorescence in situ hybridization (FISH) on brain slices from RV-injected orthotopic glioma xenografts (OGX) using markers for glutamatergic (*Slc17a6/a7*) and GABAergic neurons (*Slc32a1*). *Slc17a6* and *Slc17a7* encode for vesicular glutamate transporters 1 and 2 (VGLUT1 and VGLUT2), respectively, whereas *Slc32a1* encodes the vesicular GABA transporter (VGAT).

Among cortical GINs innervating onto striatal gliomas, approximately 76% were glutamatergic, 19% showed dual glutamatergic and GABAergic expression, and the remainder were neither (Fig. S2*C* and *D*, 3*I*). In contrast, local GINs innervating onto striatal gliomas are mainly GABAergic (Fig. S2*E* and *F*). In thalamic gliomas, 84% of cortical GINs were glutamatergic, 6% dual-expressing, and 10% neither (Fig. 3*G, H* and *I*). For motor cortex gliomas, about 85% of cortical GINs were glutamatergic, 7.7% dual-expressing, and 7.3% neither (Fig. S2*C*, 3*I*). Control analyses of nearby unlabeled neurons revealed no dual-positive cells, suggesting that dual expression may be influenced by glioma interactions or transcriptional reprogramming induced by rabies virus infection (25). However, *Slc32a1* has been reported to be downregulated in rabies-infected cortical neurons compared with noninfected cortical neurons (25).

Our study provides compelling evidence that long-range glioma-innervating neurons (GINs) are predominantly glutamatergic, offering crucial presynaptic validation to previously documented excitatory postsynaptic currents in glioma cells. Additionally, we reveal a remarkable diversity within the GIN population, including local GINs that are primarily GABAergic and a distinct subset of cortical GINs that exhibit dual glutamatergic and GABAergic markers. These results underscore the complexity of synaptic interactions in the glioma microenvironment, illustrating a heterogeneous synaptic landscape.

### Glioma-innervating neurons altered action potential waveforms

Gliomas modulate brain excitability through mechanisms such as enhanced extracellular glutamate release and reduced glutamate uptake (3, 26, 27). To explore how glioma-neuron synaptic connections affect the excitability of neurons innervating glioma sites, we conducted whole-cell current-clamp recordings of RV-infected neurons in the contralateral motor cortex of RV-injected motor cortex orthotopic glioma xenografts (OGX) (Fig. 4*A*). Control experiments utilized adenovirus (AAV) injections with a Cre-dependent TVA^TC66T^-GFP-N2cG and Cre AAV into the coordinates corresponding to the motor cortex glioma sites for three weeks, followed by RV injection and subsequent electrophysiological recordings on RV-infected neurons in the contralateral motor cortex (Fig. 4*A*). In this manner, the control cells were also trans-synaptically labeled by RV. To avoid confounding effects of RV infection, comparisons were not made directly with adjacent, non-infected neurons.

**Figure. 4.**
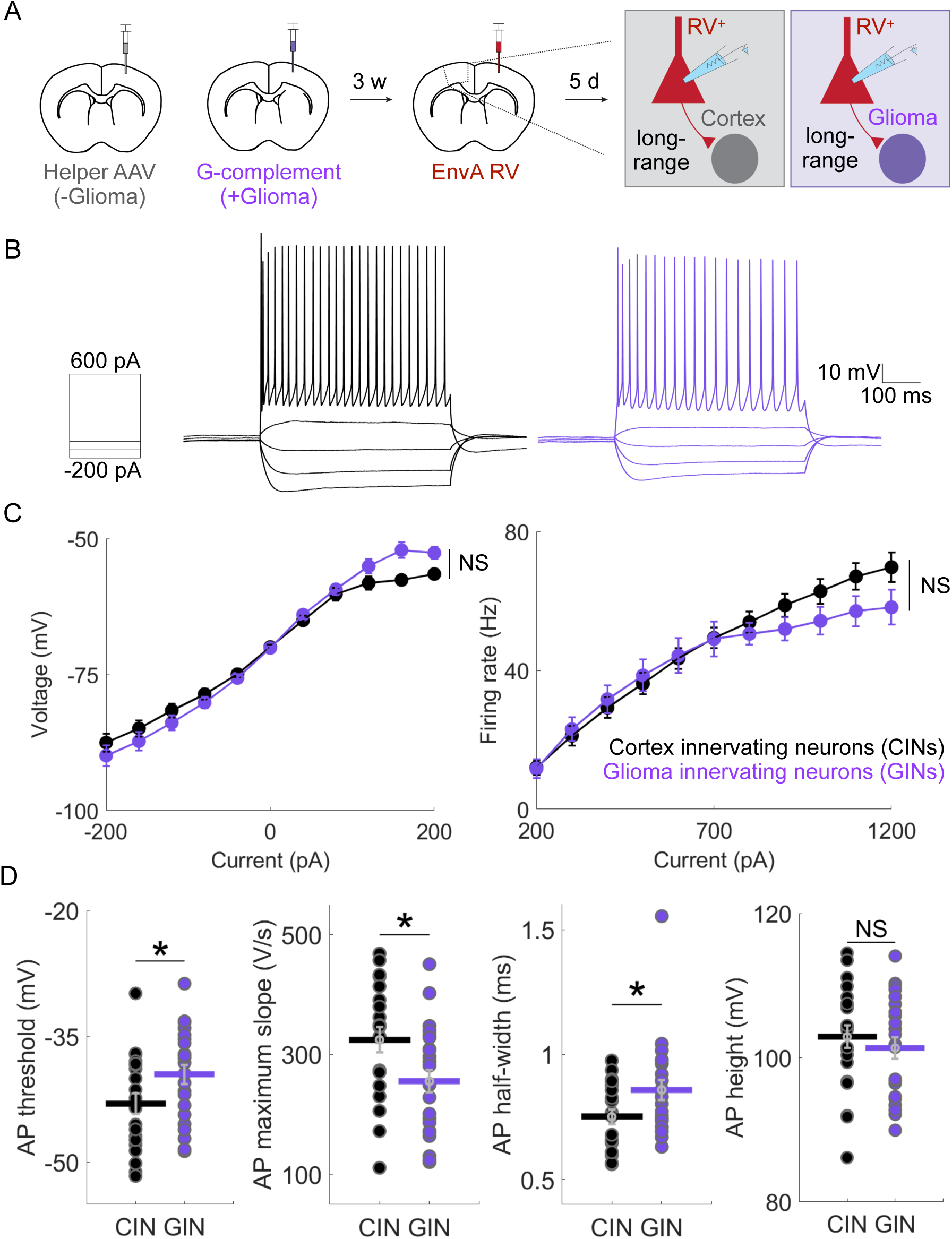
Distinct intrinsic properties of glioma-innervating neurons (GINs). (A) Schematic illustration of the experimental design for comparing intrinsic electrophysiological properties of neurons. G-complement LN229 cells (+Glioma) or helper AAVs (-Glioma, a mixture of AAV-Cre (AAV containing Cre recombinase) and AAV-FLEX-TVA^TC66T^-GFP-N2cG (AAV containing Cre-dependent expression of TVA- GFP-N2cG)) were injected into the motor cortex, followed by the administration of pseudotyped, N2cG-H2b-mCherry rabies virus (EnvA RV) for monosynaptic retrograde transsynaptic tracing of glioma-innervating neurons (GINs) or cortex-innervating neurons (CINs). Whole-cell recordings were then performed on RV-positive neurons in the contralateral motor cortex. (B) Example current-clamp recordings displaying the voltage responses of GINs (purple) and CINs (black) to a series of current injections (-200 pA, -120 pA, -40 pA, 40 pA, and 600 pA). (C) Comparative analysis of voltage (mV, *left*) and firing rate (Hz, *right*) responses between GINs and CINs, showing similar patterns (mean ± SEM displayed, n=23 GINs across 5 mice, n=21 CINs across 3 mice). NS: not statistically significant, unpaired t-tests (D) Detailed electrophysiological characterization highlights distinct differences in action potential (AP) properties. GINs exhibit a more depolarized action potential (AP) threshold, reduced maximum AP slope and increased AP half-width compared to CINs, whereas AP height remains similar across both neuron types. \**P* < 0.05, NS: not statistically significant, unpaired t-tests.

Electrophysiological analysis of 21 RV-infected neurons from 3 control animals and 23 RV-infected neurons from 5 motor cortex OGX models revealed comparable passive cellular properties and recording quality between GINs and cortex-innervating neurons (CINs): average input resistance of GINs and CINs was 127.4 ± 18.7 MOhm and 107.4 ± 13.6 MOhm; series resistance was 10.7 ± 0.8 MOhm for GINs and 11.7 ± 0.7 MOhm for CINs; membrane potential was -76.5 ± 1.5 mV for GINs and -76.9 ± 1.8 mV for CINs. Resting membrane potential and membrane time constants were also similar (Fig. 4*D*). While voltage responses and firing rates to current injections were comparable (Fig. 4*B* and *C*), action potential (AP) characteristics differed significantly (Fig. 4*D*). Specifically, GINs exhibited a more depolarized AP threshold (p=0.044), greater AP half-width (p=0.037), and reduced maximum AP slope (p=0.018) (Fig. 4*D*). These findings indicate a distinct electrophysiological signature in GINs, underlining the complex interplay between glioma cells and their neuronal inputs.

## Discussion

We established a system to map and characterize the interactions between glioma-innervating neurons (GINs) and glioma cells, offering insights into the innervation of these tumors within the brain’s unique microenvironment. By utilizing human glioma cell lines with rabies virus (RV)-based monosynaptic tracing (Fig. 1*A* and 2*A*), we demonstrated that GINs integrate broadly into pre-existing neuronal networks (Fig. 2*C*, *D, E* and 3*E*), suggesting that they do not generally favor one class of input over another. Long-range projecting cortical GINs primarily exhibit glutamatergic characteristics, with a subset also expressing dual glutamatergic/GABAergic markers, pointing to a multifaceted neuromodulatory capability (Fig. 3G, H, I, and S2*C*). Local striatal GINs, predominantly GABAergic, underscore the diversity in neurotransmitter expression depending on glioma location (Fig. S2*E* and *F*). The identification of distinct electrophysiological signatures specific to GINs suggests a potential bidirectional perturbation in the neuron-glioma circuits (Fig. 4) Our findings elucidate the complex synaptic relationships between gliomas and their neuronal inputs, offering a foundation for therapeutic strategies aimed at targeting specific neuromodulatory pathways influenced by GINs.

Human gliomas predominantly arise from the cerebral lobes (86%), most frequently in the frontal (40%), temporal (29%), parietal (14%), and occipital lobes (3%) (17). Additionally, deep structures account for 6.4%, the brainstem for 4.1%, and the cerebellum for 1.5% (17). To replicate this diversity, we selected the striatum, thalamus, and motor cortex as injection sites to represent the heterogeneity of the brain microenvironment and leverage their complex neuromodulatory pathways for our investigations. Our results suggest that gliomas integrate into existing neuronal inputs. This integration process is likely facilitated through synapses with nearby neuronal axon terminals, although the exact mechanisms by which glioma cells connect these synapses and their impacts on synaptic terminals remain to be explored.

Our comprehensive mapping of GINs revealed their localization in key cholinergic, dopaminergic, histaminergic, and serotonergic centers across the brain, suggesting that a range of neuromodulators beyond glutamate and GABA could be involved in neuron-glioma interactions and contribute to glioma growth. While further validation is necessary to confirm these findings, the implication of such diverse neuromodulatory involvement may be profound. For instance, modulating the cholinergic signaling induces cell death in glioblastoma stem cells (28). Additionally, recent studies utilizing RV retrograde tracing also highlight the important role of acetylcholine signaling in both neuron-glioma synapses and neuronal-mediated glioma growth (29)(https://doi.org/10.1101/2024.03.18.585565). Moreover, the emerging therapeutic agent dordaviprone, which targets the Dopamine receptor D2 (DRD2), has shown a 20% overall response rate in treating H3 K27M–mutant diffuse midline glioma in a phase II clinical trial, with a subsequent phase III study underway (30, 31). Elevated *DRD2* expression in clinical glioblastoma specimens and its associated pro-proliferative effects highlight the intricacies of dopaminergic signaling within glioma pathology (32). Furthermore, the expression of Dopamine receptor D5 counteracts the cytotoxic effect of DRD2 antagonism, adding a layer of complexity to the dopaminergic regulation in glioma (33). The mechanism in which dopaminergic signaling affects neuron-glioma interaction presents a promising area for future research.

The predominance of glutamatergic interactions aligns with previous observations that glutamatergic signaling induces tumor growth via an AMPAR-dependent mechanism (3, 4). The observation of GINs with dual glutamatergic/GABAergic markers added a layer of complexity, indicating that gliomas may not only receive excitatory neuronal signals but also be affected by inhibitory pathways. Furthermore, our analysis of OGX in the striatum indicates robust local innervation which, as the striatum has no glutamatergic neurons, indicates strong interactions with local GABAergic neurons. This complexity is further highlighted by a recent preprint emphasizing the role of GABA-mediated synaptic interaction and growth in diffuse midline glioma (https://doi.org/10.1101/2022.11.08.515720), suggesting that inhibitory neurotransmission plays a crucial role in glioma biology.

Transsynaptic tracing with RV has transformed our ability to study neural circuits and comprehensively map input neurons (11). Traditionally, this method has been confined largely to anatomical identifications due to its cytotoxic effects on infected neurons. In this study, we utilize a newer generation N2c rabies virus, which enhanced both the efficiency of transsynaptic labeling and neuronal viability (15). To ensure the precision of our molecular and electrophysiological analyses, we limited the infection period to five days, mitigating potential confounding effects from the virus. This approach allowed us to delineate differences in action potential (AP) characteristics between GINs and CINs. The more depolarized AP threshold, greater AP half-width and reduced AP maximal slope are consistent with a reduction in sodium channels and potential overall excitability in GINs, even if no statistically significant differences in AP firing rates were observed. Potential explanations include direct homeostatic effects from the gliomas or retrograde effects from the connecting glioma cells, such as endocannabinoid signaling or transsynaptic bridges (34–36). Further studies are needed to elucidate the mechanisms underlying these observations.

Despite our findings, the reliance on viral transduction and the use of a specific human glioma cell line in an orthotopic mouse model may not capture the full spectrum of tumor-neuron interactions present in human gliomas. Excitingly, similar distributions and molecular features of GINs were observed by independent groups using different model systems, underscoring the generalizability and relevance of our discoveries (29)(https://doi.org/10.1101/2024.03.18.585565).

Looking forward, further exploration into how glioma recruits and forms synapses with neuronal axonal terminals, beyond known mechanisms like Neuroligin 3 and BDNF, and how glioma cells benefit from other neuromodulatory signaling, will be crucial (6–8). Understanding these interactions at a deeper level could unveil new strategies for disrupting tumor-promoting neuronal signals, leading to novel therapeutic approaches to combat glioma. Future research should also extend these methodologies to other cancer types that interact with the nervous system, offering insights into how malignancies exploit host physiology across a broader spectrum.

In conclusion, our research revealed the brain-wide pattern of neuronal connections to gliomas and showed that it involves diverse neurotransmission and neuromodulatory pathways. These glioma-innervating neurons exhibit distinct molecular and electrophysiological features, providing a foundation for future studies aimed at exploring the detailed mechanisms of neuronal contributions to glioma pathology and their implications for glioma treatment strategies.

## Materials and Methods

Further information and requests for reagents may be directed to Bernardo Sabatini.

### Cloning and packaging of lentiviral constructs

The coding sequences for TVA-P2A-EGFP-P2A-N2cG (G-complement) (37) and TVA-P2A-EGFP (Control) were synthesized (Integrated DNA Technologies) and cloned by Gibson assembly into a lentiviral vector downstream of the full-length EF1α promoter. Correct assembly was verified by whole-plasmid sequencing (Plasmidsaurus). Lentivirus was produced as previously described (38). Briefly, HEK293T cells were transfected (Mirus Trans-IT) with 1.6μg of lentiviral vector, 1μg of psPAX2 (Addgene #12260), and 0.4μg of VSV-G (Addgene #8454) per well of a 6-well plate. Media was replaced after 24 hours and viral supernatant was collected 48 hours post-transfection and passaged through a 0.45μm filter.

### Cell Culture and transduction

Human glioblastoma LN-229 cell line (CRL-2611™) were purchased from ATCC. Cells were cultured in DMEM (Life Technologies Corporation; cat. #11995-065), supplemented with 10% FBS (SeraPrime; cat. #F31016-500) and 1% of penicillin-streptomycin (Thermo Fisher Scientific; cat #15140122). The cells were grown at 37°C with 5% (v/v) CO_2_. Once a plate was confluent, cells were detached with 0.05% trypsin-EDTA (Life Technologies Corporation; cat. #25300-120) and washed with Ca2+- and Mg2+-Dulbecco’s free phosphate buffered saline (DPBS) (Gibco; cat. # 14190144). Recovered cells were stored in 2ml Corning Cryogenic Vials (Sigma-Aldrich) in Bambanker Cell Freezing Media (Wako Chemicals USA Inc.; cat. # 302-14681). To establish control and G complement LN-229 cell line, cells were seeded at the density of 200k/well in 6 well plate and 400μl of control (TVA-eGFP) or G-complement (TVA-N2cG-eGFP) lentivirus-containing media were added to the cells. Cells were replated with fresh media after 48 hours.

### Fluorescence-activated Cell Sorting

Control (TVA-eGFP) or G-complement (TVA-N2cG-eGFP) were trypsinized 6 days after transduction. Cells were pelleted at 300g for 5 minutes and media was removed. Cell pellet was resuspended with fresh media and filtered through 70 μm cell strainer (Falcon^TM^; cat. # 352350). eGFP-positive cells were sorted and collected in ice cold fresh media using BD FACSAria™ II Cell Sorter in collaboration with the HMS Flow Core. Collected cells were immediately pelleted and resuspended in fresh media before passaging.

### Mice

We used an immunodeficient NCG (NOD-*Prkdc^em26Cd52^Il2rg^em26Cd22^*/NjuCrl, Charles River 572) mouse line in the study. All mice used in this study were between 5 and 12 weeks of age. Animals were kept on a 12:12 light/dark cycle under standard housing conditions. All experimental manipulations were performed in accordance with protocols approved by the Harvard Standing Committee on Animal Care, following guidelines described in the NIH Guide for the Care and Use of Laboratory Animals (21595115). All mice brain coordinates in this study are given with respect to Bregma; anterior–posterior (A/P), medial–lateral(M/L)

### Virus Preparation

EnvA pseudotyped rabies viruses (CVS-N2c-dG-H2b-mCherry, CVS-N2c-dG-mCherry and CVS-N2c-dG-TdTomato) were used at the concentration of 10^9^ to 10^10^ genome copies/mL and purchased from Janelia farm. Cre AAV (serotype AAV retrograde, addgene cat# 55636) and recombinant AAV of serotype 8 encoding FLEX-TVA^TC66T^-GFP-N2cG under the control of a hSyn promoter was used in the study at the concentration of 1013 genome copies/mL and kindly shared by Adam Granger’s laboratory at Broad Institute. Recombinant AAV of serotype PHPeB encoding hChR (H134R)-EYFP under the control of a hSyn promoter was used in the study at the concentration of 1013 genome and purchased from Addgene (Cat# 26973-PHPeB).

### Stereotaxic Intracranial Injections

All surgery was performed in aseptic conditions. Mice were anesthetized with 2 to 3% isoflurane and placed in a stereotactic frame (David Kopf Instruments). Skulls were exposed, leveled and small holes were drilled into the skull. Ten thousand Ctrl or G-complement LN229 human glioblastoma cells were injected with 1500nl total volume at a rate of 300nl/min via Hamilton syringes. Viruses were injected (300 nL for rabies virus, 150nl for retro cre AAV/ TVATC66T-GFP-N2cG AAV 1:5 mixture, 300nl for AAV-hSyn-hChR-EYFP-PHPeB) at a rate of 60 nL/min, 30nl/min or 60 nL/min via glass pipettes with tip size of ∼40 μm. Mice were given pre- and postoperative oral carprofen (MediGel CPF, 5 mg/kg/day) as an analgesic, and monitored daily for at least 4 d postsurgery. All coordinates that were used in this study were relative to Bregma (in millimeters) and were as follows: for striatum: 0.3 A/P, 2 M/L, 3.5 depth; for thalamus: -1.8 A/P, 1.5 M/L, 3.8 depth; for motor cortex: 1 A/P, 1.2 M/L, 1.5-1.7 depth, for the optogenetic experiment: 2.1 A/P, +/-0.7 M/L, 1.7 depth; 2.1 A/P, +/-1.7 M/L, 1.7 depth; -1 A/P, 1.2 M/L, 1.5 depth.

### Whole-Brain Rabies Tracing

TVA, a receptor of an avian virus envelope protein (EnvA), and N2cG were introduced through G-complement LN229 cells or AAV encoding Cre-dependent genes (AAV8-hSyn-FLEX-TVATC66T-GFP-N2cG) and retro-Cre injected unilaterally (150 nL of 5:1 mixture). After 3 wk of tumor implant or AAV expression, 300 nL of G-deleted rabies virus pseudotyped with EnvA (CVS-N2c-dG-H2b-mCherry or CVS-N2c-dG-mCherry) was injected intracranially. RV-injected animals were perfused transcardially with ice-cold phosphate-buffered saline (PBS) followed by 4% PFA, 5 d after RV injection. After a 24-h postfix in 4% PFA, brains were kept in 30% sucrose for 3-5 days and then freeze in freezing media (Tissue-Tek O.C.T.). Whole brain cryosection with cryostat (Leica, CM1950) was then performed on frozen brains and the sections were collected in PBS. Every one out of four sections were mounted with DAPI fluoromount-G (SouthernBiotech, CAT#0100-20). Slices were imaged with an Olympus VS200 slide scanning microscope or a spinning disk confocal microscope.

### NeuroInfo automatic cell detection and quantification

Coronal sections from the brain were collected in a 24-well plate, with four successive sections placed in each well. One section from each well was then mounted onto slides, with each slide containing around 12 slices on average. After the slides were scanned with an Olympus VS200 slide scanning microscope, the images were imported to NeuroInfo software (MBF Bioscience, Williston, VT) where each section was individually outlined. Starting with section 1, adjacent sections in the whole brain series were overlaid and aligned automatically, with manual adjustments made as needed. The aligned whole brain image was reconstructed into an image stack and registered to the Allen Mouse Brain Atlas. Each image from the stack was registered to the template image manually. After mapping, neurons were identified using the “cell detection” pipeline. The maximum and minimum cell size parameters were set, and the cell detection tool was applied to an image for the specific channel containing the cells to be detected (red/TRITC for mCherry-positive neurons). Detected cells were marked with selected markers. Subsequently, the cell strength threshold was adjusted to identify only mCherry-positive neurons. Various brain sections were sampled and tested with the adjusted threshold to ensure optimal parameters for detecting neuron cell bodies. After finalizing these values, the cell detection function was applied to the entire brain image stack. After cell detection was complete, markers detected false-positive cells were manually removed as needed and false-negative cells were manually added. Markers of the experiment was then mapped onto the atlas. In addition to viewing detected cells registered to the atlas, an Excel spreadsheet was generated, listing the number of mCherry-positive cells in each brain structure. This data included cells counts for each brain structure in both hemispheres and the mean cell strength value for the cells in each structure. The data from the excel spreadsheet was further extracted and analyzed using MATLAB and Graphpad. The number of total input neurons in the brain was normalized by the total number of mCherry-positive neurons.

### In Situ Hybridization

For fluorescent in situ hybridization, animals were anesthetized with isoflurane before decapitation, and the brains were rapidly removed and frozen. A total of 20 μm prefrontal and tumor injection sites (striatum, thalamus and motor cortex) coronal sections of freezing media embedded brains (Tissue-Tek O.C.T.) were prepared on cryostat (Leica, CM1950), and mounted on SuperFrost Plus glass slides (VWR). Multiplexed fluorescent in situ hybridization was performed using the ACDBio RNAScope reagents in collaboration with Harvard Medical School Neurobiology Imaging Facility. Images were acquired via spinning disk confocal microscope. Images were acquired using either a 20x or 60x oil objective on a confocal microscope (Yokogawa W1 spinning disk confocal, Nikon or LSM710, Zeiss).

### Electrophysiology

For whole-cell patch clamp recordings, mice were anesthetized by isoflurane inhalation and transcardially perfused with ice-cold artificial cerebral spinal fluid (ACSF) consisting of (in mM): 125 NaCl, 2.5 KCl, 25 NaHCO_3_, 2 CaCl_2_, 1 MgCl_2_, 1.25 NaH_2_PO_4_, and 17 glucose (300 mOsm kg^-1^). After decapitation of the mouse, the brain was removed and transferred for slicing on a Leica VT1200s vibratome. Coronal slices of 300 μm thickness were prepared in ice-cold, carbogen-saturated ACSF and incubated for ∼10 minutes in choline-based solution at 34°C consisting of (in mM): 110 choline chloride, 25 NaHCO_3_, 2.5 KCl, 7 MgCl_2_, 0.5 CaCl_2_, 1.25 NaH_2_PO_4_, 25 glucose, 11.6 ascorbic acid, and 3.1 pyruvic acid. The slices were then incubated in a second chamber containing ACSF at 34°C for 1 hour. Following this incubation, the chamber was cooled to room temperature until use. ACSF and choline solutions were continuously bubbled with carbogen gas (95% O_2_, 5% CO_2_).

Recordings were conducted in a recording chamber at 30-32°C with perfusion of carbogen-bubbled ACSF at a flow of 4-6 ml min^-1^. A Scientifica microscope equipped with a digital camera (Hamamatsu) and epifluorescence (LED light source by CoolLED) was used for visualization of rabies-mCherry^+^ neurons in primary and secondary motor cortex using a 60x/0.9NA LUMPlanFl/IR Olympus water-immersion objective. Patch pipettes (2.0 – 3.5 MOhm) were pulled from borosilicate glass (Sutter Instruments) and filled with K-based internal solution (in mM): 135 KMeSO_3_, 3 KCl, 10 HEPES, 1 EGTA, 0.1 CaCl_2_, 4 Mg-ATP, 0.3 Na-GTP, and 8 Na_2_Phosphocreatine (pH 7.3 adjusted with KOH, 290 mOsm kg^-1^). NBQX (10 μM, Tocris), CPP (10 μM, Tocris), and SR 95531 hydrobromide (10 μM, Tocris) were bath-applied. Current-clamp recordings were acquired using a Multiclamp 700B amplifier (Molecular Devices) with a filtering frequency of 10 kHz and a digital input rate of 50 kHz (acquisition board by National Instruments). During the recording, we measured responses to current steps between -200 pA and 1200 pA under the following two conditions: (1) the neuron’s endogenous resting potential and (2) a membrane potential of -70 mV maintained by the ongoing injection of current. Data collection and analysis were performed using MATLAB with custom-written script and lab software (ScanImage).

### Immunohistochemistry

Mice were anesthetized with isoflurane and perfused transcardially with PBS followed by 4% paraformaldehyde (PFA). Collected brains were post-fixed in 4% PFA overnight and transferred to 30% sucrose at 4°C. Brain tissues were then frozen in freezing media (Tissue-Tek O.C.T.) followed by cryostat sectioning in the coronal plane at 50 μm thickness (Leica CM3050 S) and stored in PBS at 4°C. The left hemisphere of the brain was slightly slit with a razor prior to sectioning to enable accurate identification of the hemisphere once the brains were sliced. Selected brain slices were washed four times with PBST (0.1%Triton X in 1x PBS) for 5 minutes each on an orbital shaker at room temperature. Brain slices were blocked with blocking solution (1% BSA and 0.02% Triton X in 1x PBS) for 2 hours on an orbital shaker at room temperature followed by incubation of primary antibodies diluted in carrier solution (3% Normal Goat/Donkey Serum [NGS/NDS] and 0.1% TritonX in 1x PBS) at 4°C overnight on a rotator. The following primary antibodies were used: guinea pig anti-NeuN (Millipore, ABN90P, 1:1000) and rabbit anti-GFAP (Agilent, Z0334, 1:500). Next day, slices were washed four times with PBST for 5 minutes each on an orbital shaker at room temperature, and then incubated with diluted secondary antibodies in carrier solution for 2 hours at room temperature. The following secondary antibodies were used: anti-guinea pig Alexa Fluor 488 (Thermo Fisher Scientific, A-11073, 1:500) and goat anti-rabbit Alexa Fluor 647 (Thermo Fisher Scientific, A-32733, 1:500). After the incubation of secondary antibodies, slices were washed four times with PBST for 5 minutes each on an orbital shaker at room temperature. Subsequently, slices were stained with DAPI diluted in 1x PBS for 20 minutes on an orbital shaker at room temperature. Finally, brain slices were washed four times with PBST for 5 minutes each and then mounted on a glass slide in mounting medium (SouthernBiotech, 0100-01), covered with glass coverslips, and sealed with nail polish. Images were acquired using either a 20x or 60x oil objective on a confocal microscope (Yokogawa W1 spinning disk confocal, Nikon or LSM710, Zeiss).

### Quantification and Statistical Analysis

Data points are stated and plotted as mean values ± SEM. P values are represented by symbols using the following code: * for 0.01 < P < 0.05, ** for 0.001 < P < 0.01, *** for 0.0001 < P <0.001, and **** for P < 0.0001. Exact P values and statistical tests are stated in the figure legends. No prior power analyses were done.

## Data Availability

All study data are included in the article and supporting information.

## ACKNOWLEDGMENTS

We thank M. EL-Rifai, E. de la Serna for their technical support; M. Albanese and A. Girasole for help with immunohistochemistry and intracranial surgery, D. Hochbaum, G. Kleinberg and G. Hulshof for help with NeuroInfo software; A. Granger, P. Capelli and D. Shi for reagent sharing; A. Granger, K. Mastro, K. Kurmi, S. Liu and members of the B.L.S. laboratory for helpful discussions and comments on the manuscript. Confocal images were acquired at the Core for Imaging Technology & Education at Harvard Medical School. Fluorescence in situ hybridization was performed by Neurobiology Imaging Facility at Harvard Medical School. This work was supported by a National Cancer Institute K12 award (K12CA090354 to A.L.H), Lubin Family Foundation Scholar Award (A.L.H), and by the NIH (M.C.H and B.L.S.).

## Author contributions

Conceptualization: A.L.H, B.L.S., M.C.H.; methodology: A.L.H., S.G., T.K.; performed research: A.L.H., S.G., T.K., M.I., W.W., Y.C., S.H, Z.L. S.J.; Analysis: A.L.H., S.G., E.A.F., Y.C.; writing (original draft), A.L.H.; writing (review and editing), A.L.H., S.G., T.K., M.I., W.W., T.C., B.L.S., M.C.H.; visualization, A.L.H., S.G., Y.C., B.L.S., M.C.H.; supervision, A.L.H., B.L.S., M.C.H.; funding acquisition, A.L.H., B.L.S., M.C.H. All authors gave feedback on the manuscript.

---

**Supplemental Figure 1.**
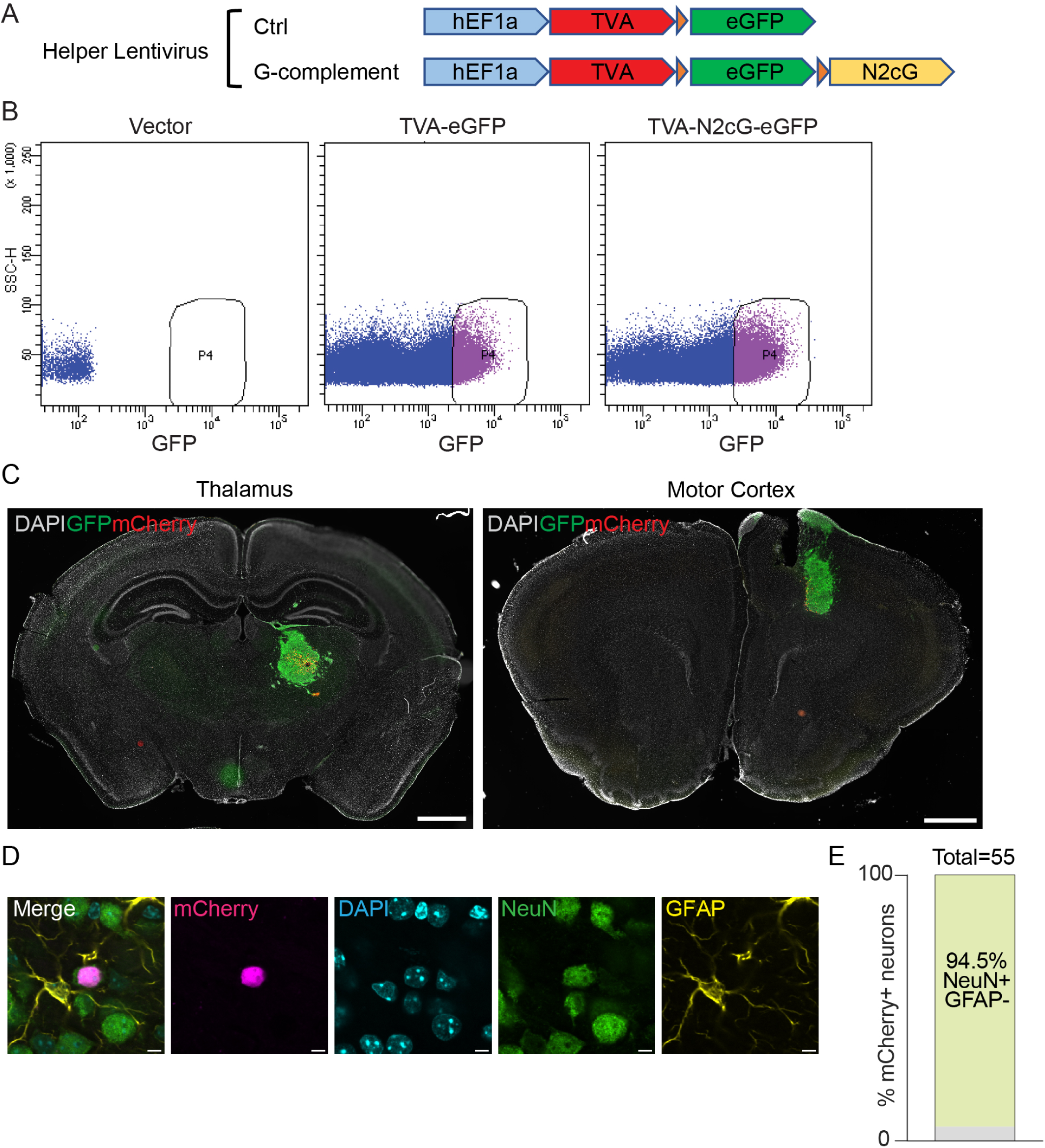
Development of glioma cell lines for rabies virus tracing and validation of specific neuronal labeling. (A) Schematic illustration detailing the construction of Ctrl and G-complement helper lentiviral plasmid used to generate LN229 glioma cell lines susceptible to rabies virus infection. (B) Flow cytometry gating for GFP positive LN229 cells transduced with TVA-eGFP or TVA-N2cG-eGFP lentivirus. Area P4 indicates the cell population collected. (C) Representative coronal brain sections from Ctrl Thalamic or Motor Cortex OGX after RV-based tracing. Images show nuclear-localized RV H2b-mCherry (red) labeling the GFP-positive OGX (starter cell population) without evidence of distant transsynaptic labeling. Scale bar: 1mm. (D) Immunohistochemistry confocal images of a representative mCherry-positive neuron from the ipsilateral cortex of striatal OGX. Cell is positively stained for neuronal nuclei (NeuN) but negative for glial fibrillary acidic protein (GFAP), indicating specific neuronal labeling. Scale bar: 5μm. (E) Quantitative bar graph displaying the percentage of mCherry-positive cells that are NeuN-positive and GFAP-negative (n=4 mice). The gray bar represents NeuN-negative and GFAP-negative.

**Figure S2.**
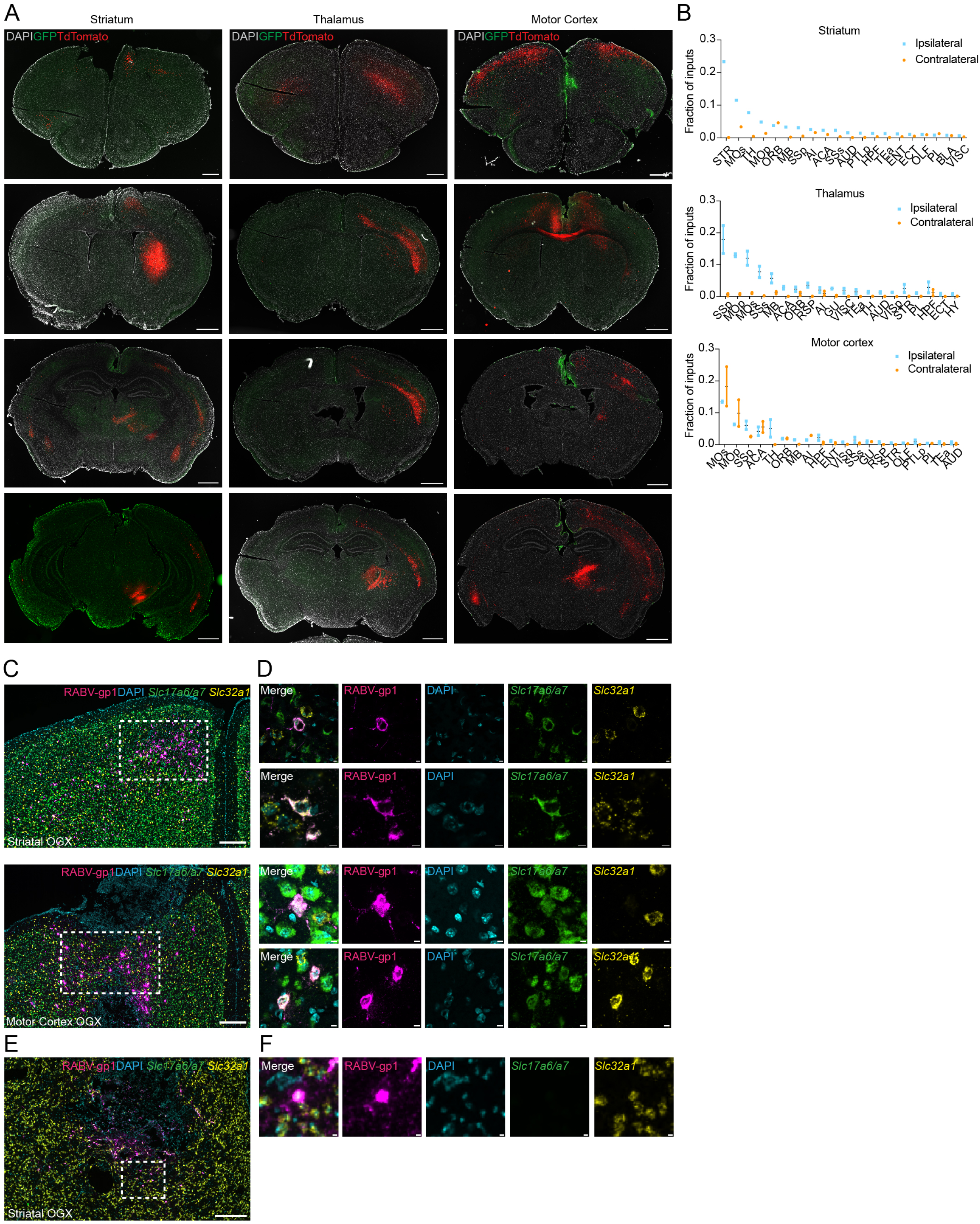
Tracing neuronal inputs in non-tumor-bearing animals with nonpseudotyped rabies virus and molecular characterization of glioma-innervating neurons. (A) Representative coronal brain sections from animals injected with nonpseudotyped, N2cG-deleted rabies virus (CVS-N2c-dG-TdTomato) at stereotaxic coordinates corresponding to glioma sites (striatum, thalamus and motor cortex) but without tumor implantation. Images display TdTomato-positive neurons and their processes, indicating whole-cell labeling. Scale bar: 500μm for the first row and 1mm for subsequent rows. (B) Top 20 input regions to the striatum, thalamus and motor cortex in non-tumor-bearing animals, showing the distribution of TdTomato-labeled neurons ipsilateral and contralateral to the injection sites (n=2 mice for the thalamus and motor cortex and n=1 mouse for the striatum; STR, striatum; MOs, secondary motor area; TH, thalamus; MOp, primary motor area; ORB, orbital area; MB, midbrain; SSp, primary somatosensory area; AI, agranular insular area; ACA, anterior cingulate area; PTLp, posterior parietal association areas; HPF, hippocampal formation; TEa, temporal association areas; ENT, entorhinal area; ECT, ectorhinal area; OLF, olfactory areas; PL, prelimbic area; BLA, basolateral amygdalar nucleus; VISC, visceral area; SSs, supplemental somatosensory area; RSP, retrosplenial area; GU, gustatory areas; VISp, primary visual area; HY, hypothalamus). (C) Representative in situ hybridization confocal image from the ipsilateral motor cortex of striatal and motor cortex OGX. RABVgp1(rabies virus complete genome)-positive cells were classified as VGLUT1/2+ (*Slc17a6 or Slc17a7 positive*)/VGAT+(*Slc32a1 positive*), VGLUT1/2+/VGAT- or VGLUT1/2-/VGAT- (no VGLUT1/2-/VGAT+ cells observed in cortex). Scale bar: 250μm. (D) Inset of panel (C) illustrating individual cells, showing VGLUT1/2+/VGAT-cell (top) and VGLUT1/2+/VGAT+ cell (bottom). Scale bar: 5 μm. (E) Representative in situ hybridization image from the ipsilateral striatum of striatal OGX. RABV-gp1-positive cells were mostly VGLUT1/2-/VGAT+. Scale Bar: 200 μm. (F) Inset of panel (E) illustrating an individual cell, showing VGLUT1/2-/VGAT+ cell. Scale bar: 5 μm.

## References

1. E. Jung et al., Emerging intersections between neuroscience and glioma biology. Nature neuroscience 22, 1951–1960 (2019).

2. F. Winkler et al., Cancer neuroscience: State of the field, emerging directions. Cell 186, 1689–1707 (2023).

3. H. S. Venkatesh et al., Electrical and synaptic integration of glioma into neural circuits. Nature 573, 539–545 (2019).

4. V. Venkataramani et al., Glutamatergic synaptic input to glioma cells drives brain tumour progression. Nature 573, 532–538 (2019).

5. V. Venkataramani et al., Glioblastoma hijacks neuronal mechanisms for brain invasion. Cell 185, 2899–2917 e2831 (2022).

6. H. S. Venkatesh et al., Neuronal Activity Promotes Glioma Growth through Neuroligin-3 Secretion. Cell 161, 803–816 (2015).

7. H. S. Venkatesh et al., Targeting neuronal activity-regulated neuroligin-3 dependency in high-grade glioma. Nature 549, 533–537 (2017).

8. K. R. Taylor et al., Glioma synapses recruit mechanisms of adaptive plasticity. Nature 623, 366–374 (2023).

9. E. Huang-Hobbs et al., Remote neuronal activity drives glioma progression through SEMA4F. Nature 619, 844–850 (2023).

10. S. Krishna et al., Glioblastoma remodelling of human neural circuits decreases survival. Nature 617, 599–607 (2023).

11. E. M. Callaway, L. Luo, Monosynaptic Circuit Tracing with Glycoprotein-Deleted Rabies Viruses. The Journal of neuroscience : the official journal of the Society for Neuroscience 35, 8979–8985 (2015).

12. I. R. Wickersham, S. Finke, K. K. Conzelmann, E. M. Callaway, Retrograde neuronal tracing with a deletion-mutant rabies virus. Nat Methods 4, 47–49 (2007).

13. M. Hyun et al., Social isolation uncovers a circuit underlying context-dependent territory-covering micturition. Proceedings of the National Academy of Sciences of the United States of America 118 (2021).

14. C. W. Mount, B. Yalcin, K. Cunliffe-Koehler, S. Sundaresh, M. Monje, Monosynaptic tracing maps brain-wide afferent oligodendrocyte precursor cell connectivity. eLife 8 (2019).

15. T. R. Reardon et al., Rabies Virus CVS-N2c(DeltaG) Strain Enhances Retrograde Synaptic Transfer and Neuronal Viability. Neuron 89, 711–724 (2016).

16. N. R. Wall, I. R. Wickersham, A. Cetin, M. De La Parra, E. M. Callaway, Monosynaptic circuit tracing in vivo through Cre-dependent targeting and complementation of modified rabies virus. Proceedings of the National Academy of Sciences of the United States of America 107, 21848–21853 (2010).

17. S. Larjavaara et al., Incidence of gliomas by anatomic location. Neuro Oncol 9, 319–325 (2007).

18. A. M. Thomson, Neocortical layer 6, a review. Front Neuroanat 4, 13 (2010).

19. J. Cox, I. B. Wicen, Striatal circuits for reward learning and decision-making. Nature reviews. Neuroscience 20, 482–494 (2019).

20. C. Varela, Thalamic neuromodulation and its implications for executive networks. Front Neural Circuits 8, 69 (2014).

21. C. Vitrac, M. Benoit-Marand, Monoaminergic Modulation of Motor Cortex Function. Front Neural Circuits 11, 72 (2017).

22. A. Venkataraman et al., Incerto-thalamic modulation of fear via GABA and dopamine. Neuropsychopharmacology : official publica=on of the American College of Neuropsychopharmacology 46, 1658–1668 (2021).

23. J. Lu, T. C. Jhou, C. B. Saper, Identification of wake-active dopaminergic neurons in the ventral periaqueductal gray macer. The Journal of neuroscience : the official journal of the Society for Neuroscience 26, 193–202 (2006).

24. M. A. Sanchez-Gonzalez, M. A. Garcia-Cabezas, B. Rico, C. Cavada, The primate thalamus is a key target for brain dopamine. The Journal of neuroscience : the official journal of the Society for Neuroscience 25, 6076–6083 (2005).

25. M. Patino et al., Single-cell transcriptomic classification of rabies-infected cortical neurons. Proceedings of the National Academy of Sciences of the United States of America 119, e2203677119 (2022).

26. Z. C. Ye, J. D. Rothstein, H. Sontheimer, Compromised glutamate transport in human glioma cells: reduction-mislocalization of sodium-dependent glutamate transporters and enhanced activity of cystine-glutamate exchange. The Journal of neuroscience : the official journal of the Society for Neuroscience 19, 10767–10777 (1999).

27. Z. C. Ye, H. Sontheimer, Glioma cells release excitotoxic concentrations of glutamate. Cancer research 59, 4383–4391 (1999).

28. I. Cristofaro et al., Activation of M2 muscarinic acetylcholine receptors by a hybrid agonist enhances cytotoxic effects in GB7 glioblastoma cancer stem cells. Neurochem Int 118, 52–60 (2018).

29. Y. Sun et al., Brain-wide neuronal circuit connectome of human glioblastoma. bioRxiv 10.1101/2024.03.01.583047 (2024).

30. I. Arrillaga-Romany et al., ONC201 (Dordaviprone) in Recurrent H3 K27M-Mutant Diffuse Midline Glioma. J Clin Oncol 42, 1542–1552 (2024).

31. I. Arrillaga-Romany et al., ACTION: a randomized phase 3 study of ONC201 (dordaviprone) in patients with newly diagnosed H3 K27M-mutant diffuse glioma. Neuro Oncol 26, S173–S181 (2024).

32. J. Li et al., Genome-wide shRNA screen revealed integrated mitogenic signaling between dopamine receptor D2 (DRD2) and epidermal growth factor receptor (EGFR) in glioblastoma. Oncotarget 5, 882–893 (2014).

33. V. V. Prabhu et al., Dopamine Receptor D5 is a Modulator of Tumor Response to Dopamine Receptor D2 Antagonism. Clinical cancer research : an official journal of the American Associa=on for Cancer Research 25, 2305–2313 (2019).

34. P. E. Castillo, T. J. Younts, A. E. Chavez, Y. Hashimotodani, Endocannabinoid signaling and synaptic function. Neuron 76, 70–81 (2012).

35. C. Costas-Insua, M. Guzman, Endocannabinoid signaling in glioma. Glia 71, 127–138 (2023).

36. S. K. Godavarthi et al., Postsynaptic receptors regulate presynaptic transmicer stability through transsynaptic bridges. Proceedings of the Na=onal Academy of Sciences of the United States of America 121, e2318041121 (2024).

37. G. Pouchelon et al., The organization and development of cortical interneuron presynaptic circuits are area specific. Cell reports 37, 109993 (2021).

38. M. J. Dolan et al., Exposure of iPSC-derived human microglia to brain substrates enables the generation and manipulation of diverse transcriptional states in vitro. Nature immunology 24, 1382–1390 (2023).

